# Exploring the impact of sequence context on errors in SNP genotype calling with Whole Genome Sequencing data using AI-based autoencoder approach

**DOI:** 10.1101/2024.03.23.586433

**Authors:** Krzysztof Kotlarz, Magda Mielczarek, Przemysław Biecek, Bernt Guldbrandtsen, Joanna Szyda

## Abstract

A critical step in the analysis of WGS data is variant calling. Despite its importance, variant calling is prone to errors. Our study investigated the association between incorrect SNP and variant quality metrics and nucleotide context. In our study, incorrect SNPs were defined in twenty Holstein-Friesian cows by comparing their SNPs genotypes identified by whole genome sequencing with the IlluminaNovaSeq6000 and the EuroGMD50K genotyping microarray. The data set was divided into the correct set of SNPs (666,333 SNPs) and the incorrect set of SNPs (4,557 SNPs). The training data set consisted of only the correct SNPs, while the test data set contained a balanced mix of all the incorrectly and correctly called SNPs. An autoencoder was constructed to identify systematically incorrect SNPs that were marked as outliers by a one-class support vector machine and isolation forest algorithms. The results showed that 59.53% (±0.39%) of the incorrect SNPs had systematic patterns, with the remainder being random errors. The frequent occurrence of the CGC trimer was due to mislabeling a call for C. Incorrect T instead A call was associated with the presence of T in the neighboring downstream position. These errors may arise due to the fluorescence patterns of nucleotide labelling.

## INTRODUCTION

The field of genomics has undergone a revolution thanks to whole-generation sequencing (WGS), which makes it possible to generate large amounts of genomic data quickly and affordably (1). A critical step in the analysis of WGS data is variant calling, which involves comparing sequenced reads with a reference genome to identify polymorphic sites, e.g., single nucleotide polymorphisms (SNP) or structural variants (SV). However, variant calling is challenging. Errors can occur in each step of variant identification, including sequencing errors, incorrect alignment to the reference genome (2), or erroneous variant calling (3).

We explored the very first step leading to variant detection, i.e. the generation of short-sequenced reads. We focussed on how the sequence context, defined as nucleotides immediately preceding or following a variant, affects the performance of variant calling, by provoking the reading of an erroneous SNP genotype (4). Although the aim appears to be straightforward, to provide a meaningful analysis of the phenomenon, one needs to address two issues, both related to the fact that fortunately the error rate in short sequence reads generated by the sequencing by synthesis workflows is quite low, ranging between 1 error per 1000 bp for the Illumina Hiseq2000 platform to 1 error per 163 bp for the Illumina MiniSeq platform (4, 5). First, for a supervised learning approach, such as all Artificial Intelligence (AI) applications, an informative and possibly large training data set is required, which in view of the low rate of wrongly called SNPs is difficult to obtain. Second, by classification of correctly and incorrectly called SNP genotypes, one encounters the problem of extreme class imbalance, with the correct SNP class being much more highly represented than the incorrect class.

Problems in the acquisition of the informative training data are typically mitigated by generating artificially incorrect variants, as e.g. in Krachunov et al. (6). However, this approach does not retain the biological context of the sequencing errors. Problems related to modelling such rare event data have been known and addressed in statistics for decades, especially that, like in the case of incorrect SNP detection, it is mostly the under-represented class that is of primary interest. Extremely imbalanced class frequency in model-based classification, e.g. in logistic regression, may result in low accuracy of classification as the under-represented class does not provide enough information for accurate estimation of model parameters. This results in classification bias towards the high-frequency class (7, 8). In logistic regression, the most frequently used approach is to weight or penalise the underlying (log)likelihood function (9, 10) or to apply stochastic techniques such as the bootstrap or Markov chain Monte Carlo (11). Also, in model-free classification, such as traditional AI implementing deep learning (DL) architectures based on neural networks, feature selection and then following feature importance estimation are biased towards the correct classification of the majority class (10). In AI, typical approaches to mitigate the problem are mostly related to input data manipulation through oversampling of the minority class or undersampling of the dominant class, as well as to the differential weighting of the contribution of the observations representing the majority and the minority class to the likelihood function.

In our study, the above-mentioned problems were addressed as follows. Training data with correct and incorrect SNP labelling and the retention of sequence context was defined by comparing genotypes identified from whole genome sequence data with the corresponding genotypes identified from the Illumina SNP oligonucleotide microarrays, which was considered as the golden standard of true genotypes. The class imbalance problem was addressed by applying an AI-based anomaly detection approach implemented as an autoencoder (AE) - a set of feed-forward neural networks that replicate input values to output values. Such an AE architecture contains a hidden core layer with fewer neurons than the input layers, which extracts the most essential set of explanatory variables that are characteristic of the input. This unsupervised learning algorithm is trained on data that in the case of our study consists of only the class of correctly called SNPs, to accurately reconstruct their genotypes in the output layer (12), which involves only training on the majority class, thereby circumventing the rare event problem represented by incorrectly called SNP class. The idea behind the detection of anomalous observations (incorrectly called SNPs) is that they do not fit the reconstruction model and consequently are identified by a high reconstruction error (13, 14).

## MATERIAL AND METHODS

### Sequenced individuals and their genomic information

Whole genome sequences and array-based SNP genotypes of twenty unrelated (no common dams and sires) Holstein-Friesian cows from a single herd were available for analysis. The cows were kept in the same barn, under identical conditions and on the same diet. The WGS were obtained from the Illumina NovaSeq 6000 platform in the paired-end mode with a 150 bp read length. The number of raw reads per individual ranged from 311,675,740 to 908,861,126, resulting in a genome-averaged coverage varying between 14X and 37X. In addition to WGS, for each cow, SNP genotypes were determined using the EuroG MD 50K genotyping platform (version 2). This is the Illumina genotyping microarray customised for the EuroGenomics Cooperative (www.eurogenomics.com), containing 52,788 SNPs.

### Identification of SNPs incorrectly called in whole genome sequences

From the WGS data, the SNPs were identified by incorporating the following workflow: (i) the quality control of the raw reads was carried out for each sample using the FastQC and MultiQC tools (15, 16), (ii) data cleaning was performed with Trimmomatic software (17) to remove low quality reads and adapter sequences, (iii) the alignment to the ARS-UCD1.2 reference genome (NCBI accession number: PRJNA391427) was performed using BWA-MEM (18) (iv) the post-alignment that included converting SAM files to binary (BAM) format, data indexing, further quality control, and marking of PCR duplicates was done by using SAMtools (19) and BEDtools (20); (v) the base quality score recalibration (BQSR), which adjusts the base quality scores of sequencing reads and (vi) the actual SNP calling was done using BaseRecalibrator and HaplotypeCaller from the GATK package (21). The data from the genotyping array was pre-processed by removing SNPs with a call rate lower than 95% and a minor allele frequency below 5%. SNPs incorrectly called by the WGS processing pipeline were identified by comparing their genotypes with those obtained from the genotyping array. Incorrectly genotyped SNPs were defined as SNPs with mismatch(es) that involved at least one allele between genotypes reported on the array and genotypes reported by the SNP calling workflow described above. Consequently, correctly called SNPs refer to full agreement in the genotypes identified by both technologies.

### Construction and training of the autoencoder

The following explanatory variables from a Variant Call Format (VCF) file resulting from the SNP calling workflow, were considered as features for the reconstruction of the SNP genotypes in the autoencoder: (a) SNP specific variables - reference allele (REF), alternative allele (ALT), (b) SNP and individual specific variables - SNP allele “A” (GT:A), SNP allele “B” (GT:B), sequencing depth of reads supporting allele “A” (AD:A), sequencing depth of reads supporting allele “B” (AD:B), and SNP genotype quality (GQ) expressed as the probability that the called genotype is true, where “A” and “B” denote two alleles of the unphased SNP genotype called for a given individual. Furthermore, the sequence context surrounding the SNP site was represented by four reference nucleotides downstream (Downstream1:Downstream4) and four reference nucleotides upstream of the SNP (Upstream1:Upstream4). All explanatory variables were transformed into principal components (PCs) by applying the PCA for the mixed data approach implemented in FactoMineR (22) in R. Various dimensionalities of data representation, expressed by different numbers of PCs, were considered.

The autoencoder algorithm was implemented in Python via the Keras interface (23) with the TensorFlow library (24). In our study, three AE architectures of increasing complexity expressed by the number of hidden layers and the number of neurons inside each layer were implemented: the small model (S) that fits only two encoding and two decoding layers, the moderate model (M) with three layers, and the large model (L) that incorporates five layers (Figure 1).

**Figure 1.**
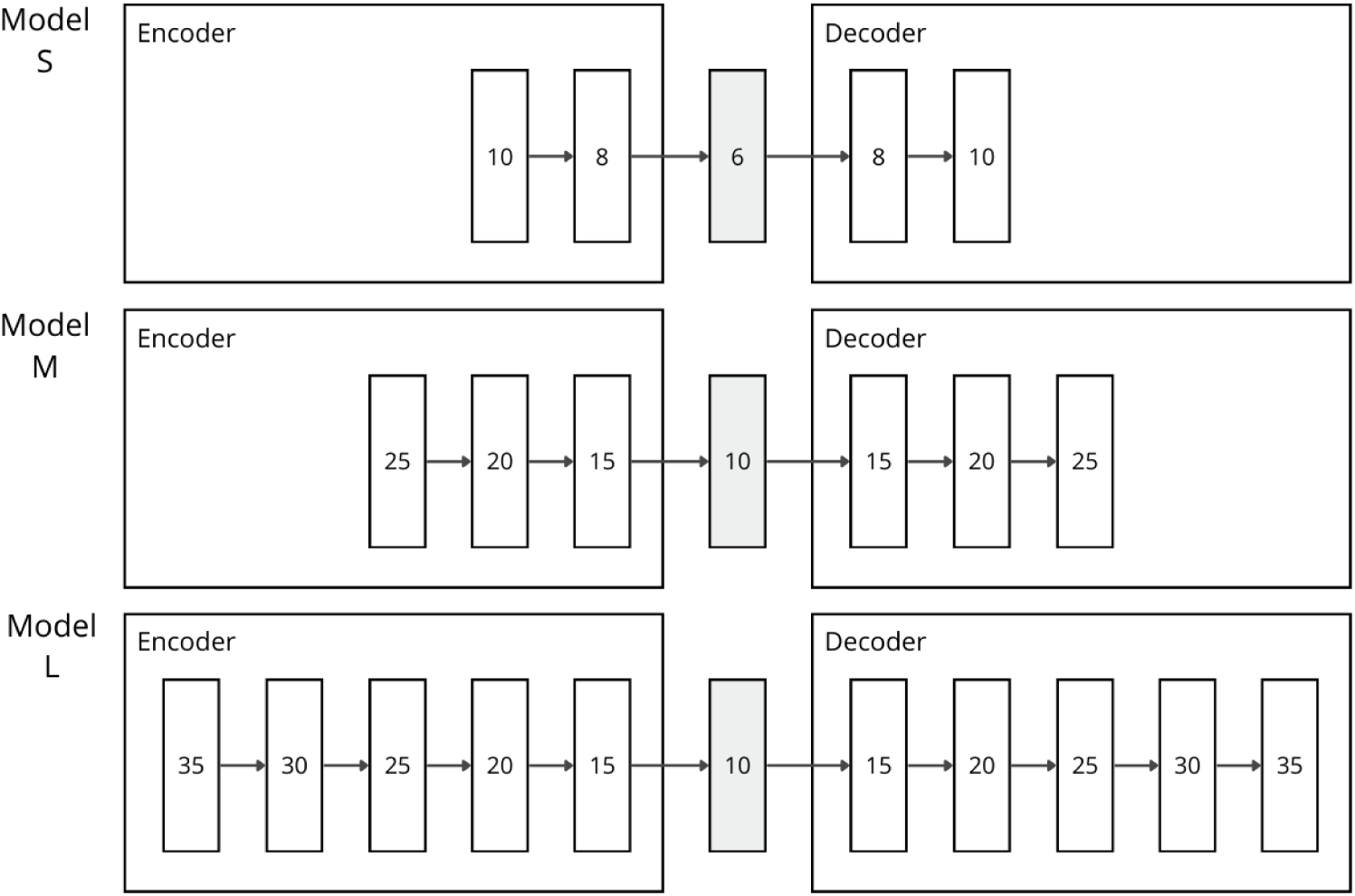
Graphical representation of autoencoder architectures (S, M, and L) implemented in our study, featuring layer configurations and the numbers of neurons.

In each model, the activation function in the layers was set to the Exponential Linear Unit (ELU) to allow for the negative input values (25), while the linear activation function was used in the last layer. The Adam optimiser (26) that implements the stochastic gradient descent algorithm with a learning rate set to 4.0۰10-3 was applied to minimise the objective (loss) function defined as the mean square error: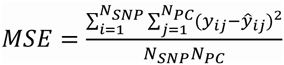 where *N_SNP_* represents the number of SNPs, *N_PC_* is the number of PCs considered, *y_ij_* represents the observed value of the j-th PC for the i-th SNP, while *ŷ_ij_* is the corresponding PC value predicted by the model.

### Data splitting and evaluation criteria

The data set was divided into (i) the set of correct SNPs containing 666,333 SNPs and (ii) the set of incorrect SNPs containing 4,557 SNPs. Incorrect SNPs made up only 0.007% of all SNPs. The training data set consisted of 661,776 SNPs that were correctly called in the WGS, while the test data set was selected as the balanced set consisting of all 4,557 incorrectly called SNPs and 4,557 correctly called SNPs that were randomly selected from the complete correctly called SNP set (Figure 2).

**Figure 2.**
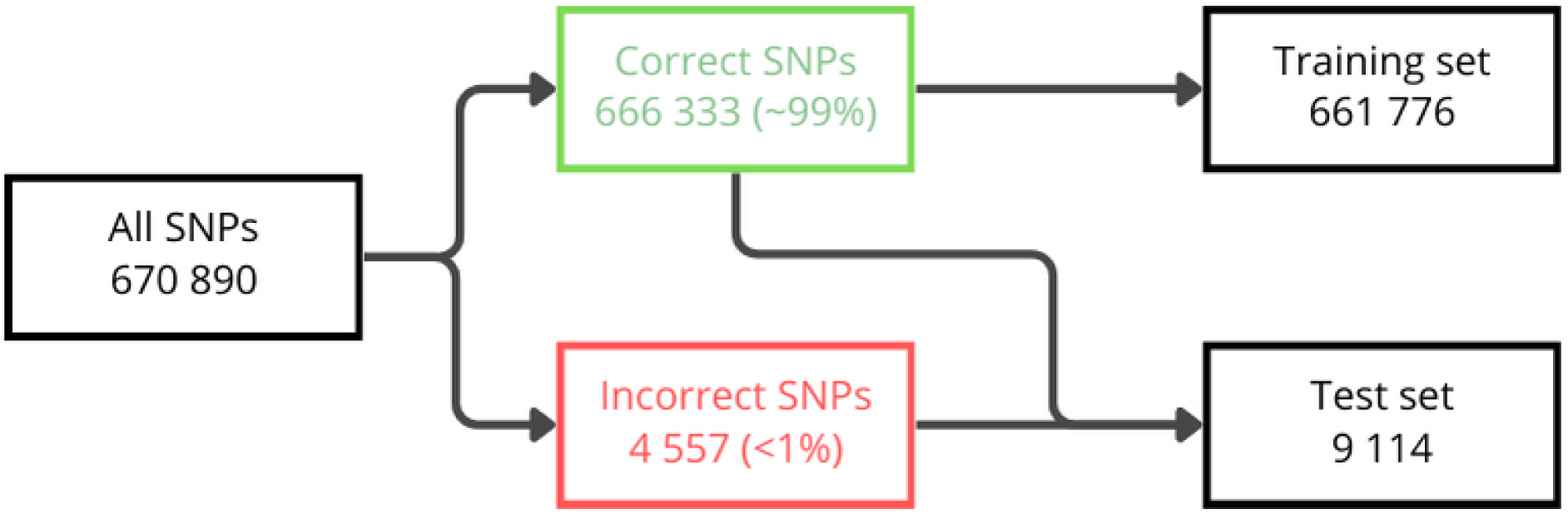
The composition of the test and the training datasets.

For the validation of the autoencoder algorithm, the 5-fold cross-validation was applied by randomly selecting 80% of the training SNPs, while the remaining SNPs formed a holdout set. The quality of reconstruction of the principal components underlying each SNP from the test data set was quantified by residuals: 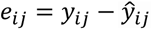 and by the Mean Absolute Error (MAE), which is given by 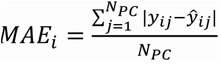, where all the elements are as defined above.

The models were trained and evaluated on an NVIDIA Tesla P100 GPU with 128 GB of RAM.

### Identification of incorrectly called SNPs

SNPs with genotypes incorrectly called in the WGS data are expected not to fit the decoding model trained by the AE on correctly called SNPs. Technically, such incorrectly called SNPs are then expected to be outliers in the reconstruction, associated with a high reconstruction error expressed by *e_ij_*. The identification of such outlying SNPs was carried out within the multidimensional space defined by PCs by applying two unsupervised learning methods. The one-class support vector machine (OCSVM) method was implemented by the scikit-learn library (27). This algorithm estimates a decision boundary that separates correctly called SNPs characterised by low reconstruction errors from incorrectly called SNPs with high reconstruction errors that are represented as outliers and are located outside the decision boundary (28). The other approach to outlier detection was the isolation forest algorithm (iForest), where each tree randomly selects a subset of PCs and creates splits until each SNP is isolated in a separate leaf node. Short paths in the decision trees represent outliers, i.e., the incorrectly called SNPs (29). In both algorithms, the maximal proportion of outliers was set to 50% to express the structure of the test data set.

The validation of the outlier detection procedure was carried out by 5-fold cross-validation on the test dataset, to select optimal values for hyperparameters for the OCSVM and iForest algorithms. Moreover, for the outlier detection, the precision metric was used to express each model’s ability to select truly incorrectly called SNPs as 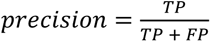, where the TP (true positive) class contains incorrectly called SNPs classified as incorrect, FP (false positive) class contains correctly called SNPs classified as incorrect, TN (true negative) class contains incorrectly called SNPs classified as correct, and FN (false negative) class contains correctly called SNPs classified as incorrect.

### Evaluation of the impact of sequence context on the systematic incorrect SNP calls

In the last part, the subset of incorrectly called SNPs was divided into systematic and random incorrect calls. Since the latter (random) incorrect calls do not associate with a specific pattern of explanatory variables, they were not further considered. On the other hand, the systematically incorrect calls were subjected to further exploration. In particular, the best classifying model, defined by the interplay between the precision metric and MAE, was then used to separate systematic SNP call errors, which in our study were represented by the TP class, from random SNP call errors represented by the TN class. The sequence context underlying the systematically incorrect SNPs was evaluated by assessing the influence of neighbouring nucleotides on the incorrect SNP call, considering the nucleotide composition on the ARS-UCD1.2 reference assembly. For this purpose, we first examined the frequencies of the nucleotides (A, C, G, T) at the four genomic positions downstream and upstream of the incorrectly called SNP. Furthermore, K-mer (K=3) frequencies composed of two bases downstream of the incorrect SNP and the alternative allele at the SNP site were calculated. The error frequency of each trimer from the TP class was normalised by its genomic frequency from all classes.

## RESULTS

### Data representation

To identify the optimal number of PCs, 11 sets starting from three PCs that accounted for 19.93% of the total variance, up to 23 PCs that jointly explained 79.07% of the total variance with an increment of two were considered (Figure 3). The contribution of each feature to consecutive PCs was shown in Figure 4.

**Figure 3.**
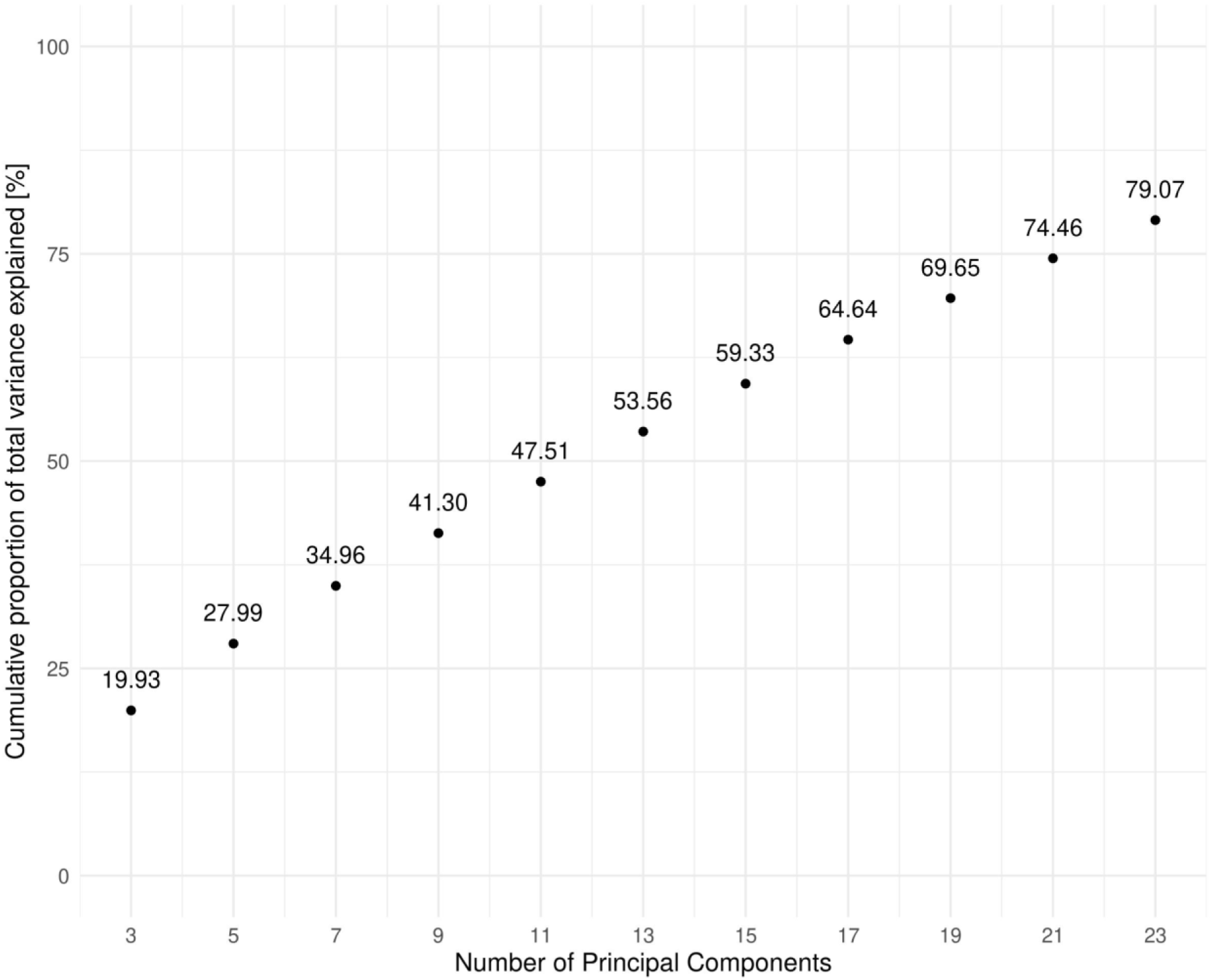
The numbers of Principal Components (PCs) used as data representation and the proportion of the total variance of variables explained by them.

**Figure 4.**
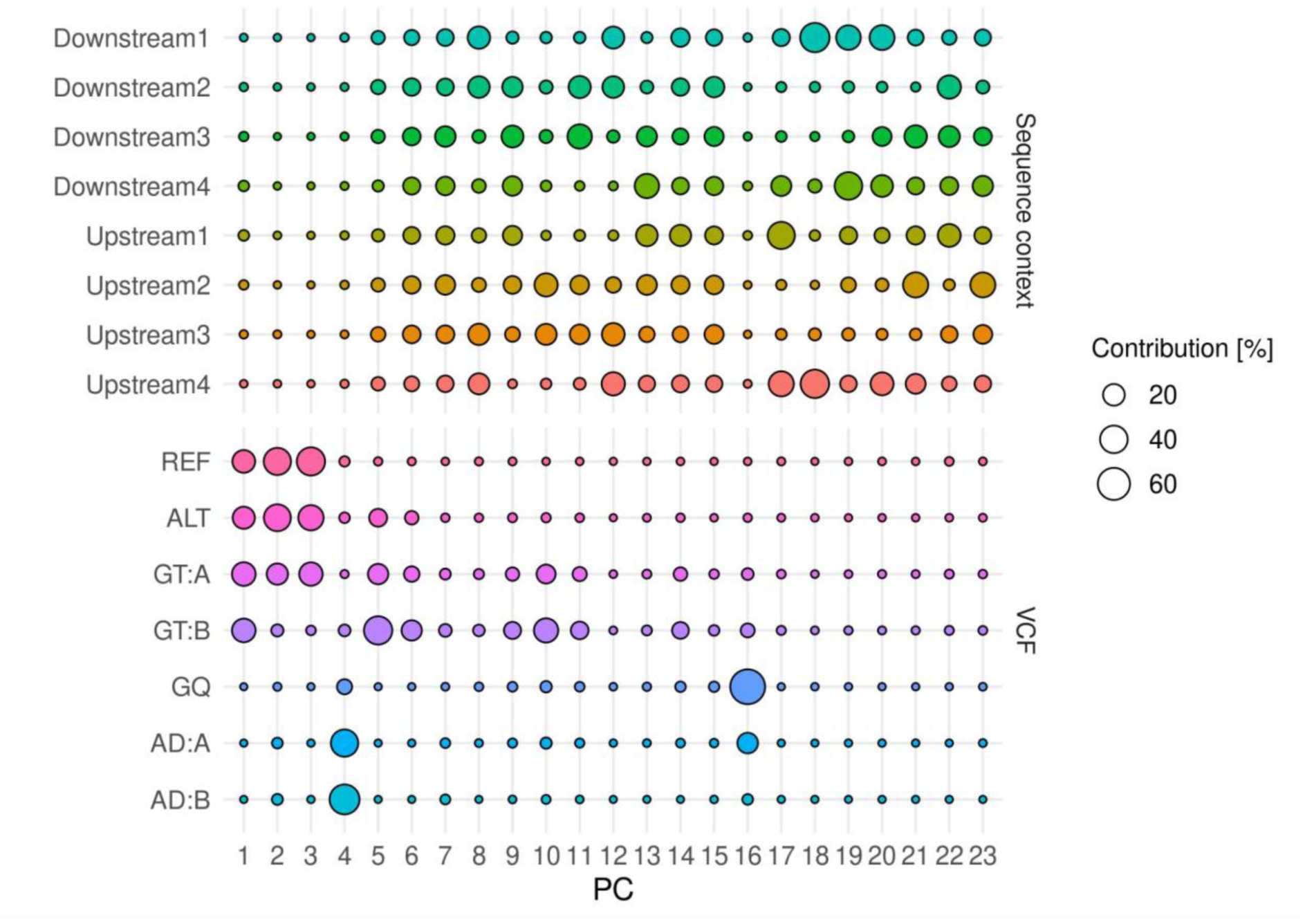
The contribution of each feature to consecutive Principal Components (PCs).

### Selection of the best model

MAEs averaged over all SNPs and all 11 results from 5-fold cross-validation of the training data set, separately for each model architecture, were shown in Figure 5. A clear pattern shows that fitting a more complex model resulted in a reduced reconstruction error.

**Figure 5.**
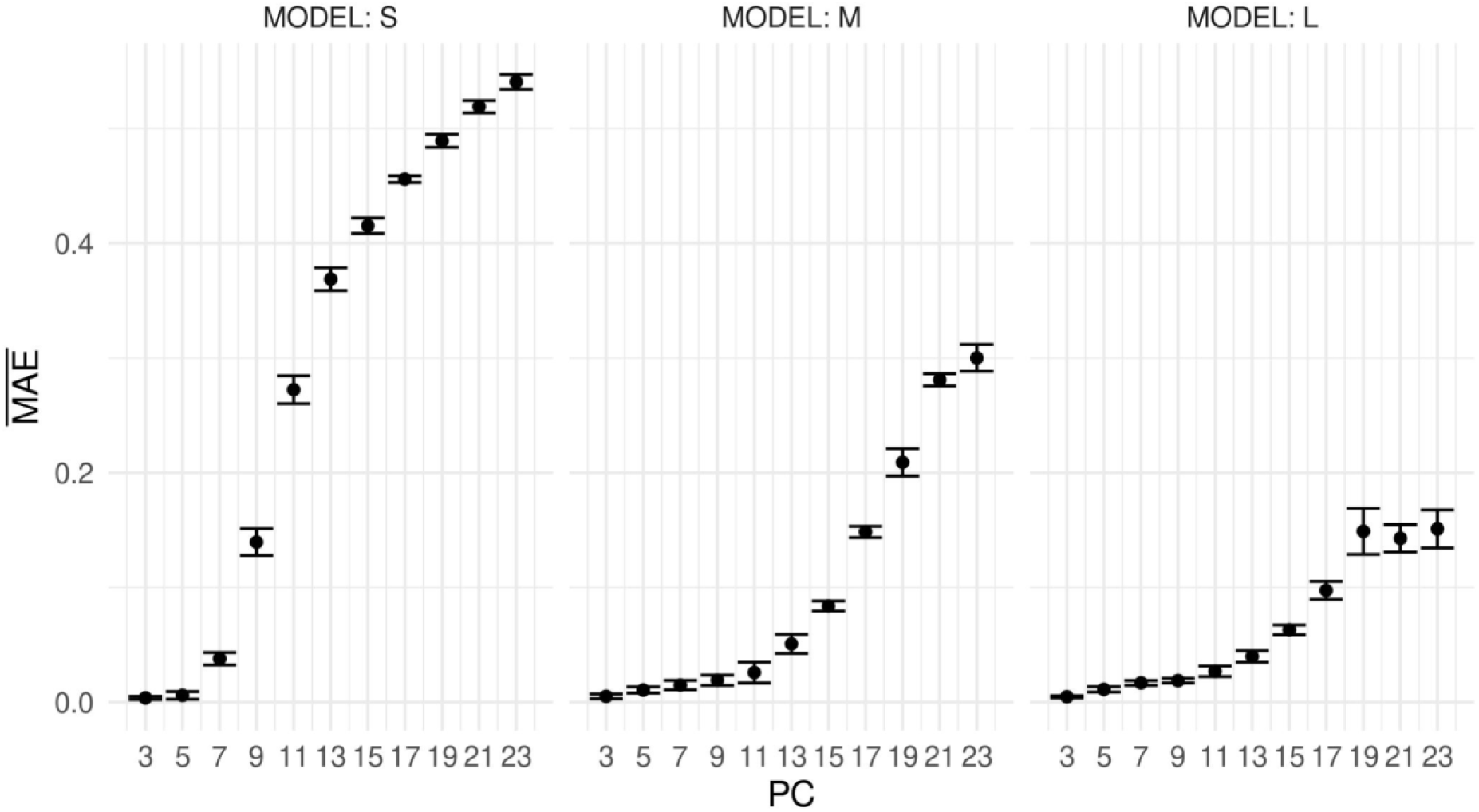
MAE averaged over all SNPs and all validation datasets derived from 5-fold cross-validation of the training data. S, M, and L represent model architectures of increasing depth and complexity. The error bars represent the 95% confidence intervals.

In the test data set, residuals corresponding to particular PCs were used to classify the SNPs as correctly called or as outliers characterised by high reconstruction error. The precision metric of the models, visualised in Figure 6, varied from 50.44% (±0.93%) in the S model that fits 5 PCs followed by the classification with iForest, to 59.53% (±0.39%) in the L model with 23 PCs followed by the OCSVM classification. Regardless of the applied classification method, the most complex model architecture, L, resulted in the highest precision. Generally, more complex model architectures that fit a larger number of PCs yielded a higher precision of classification up to 17 PCs.

**Figure 6.**
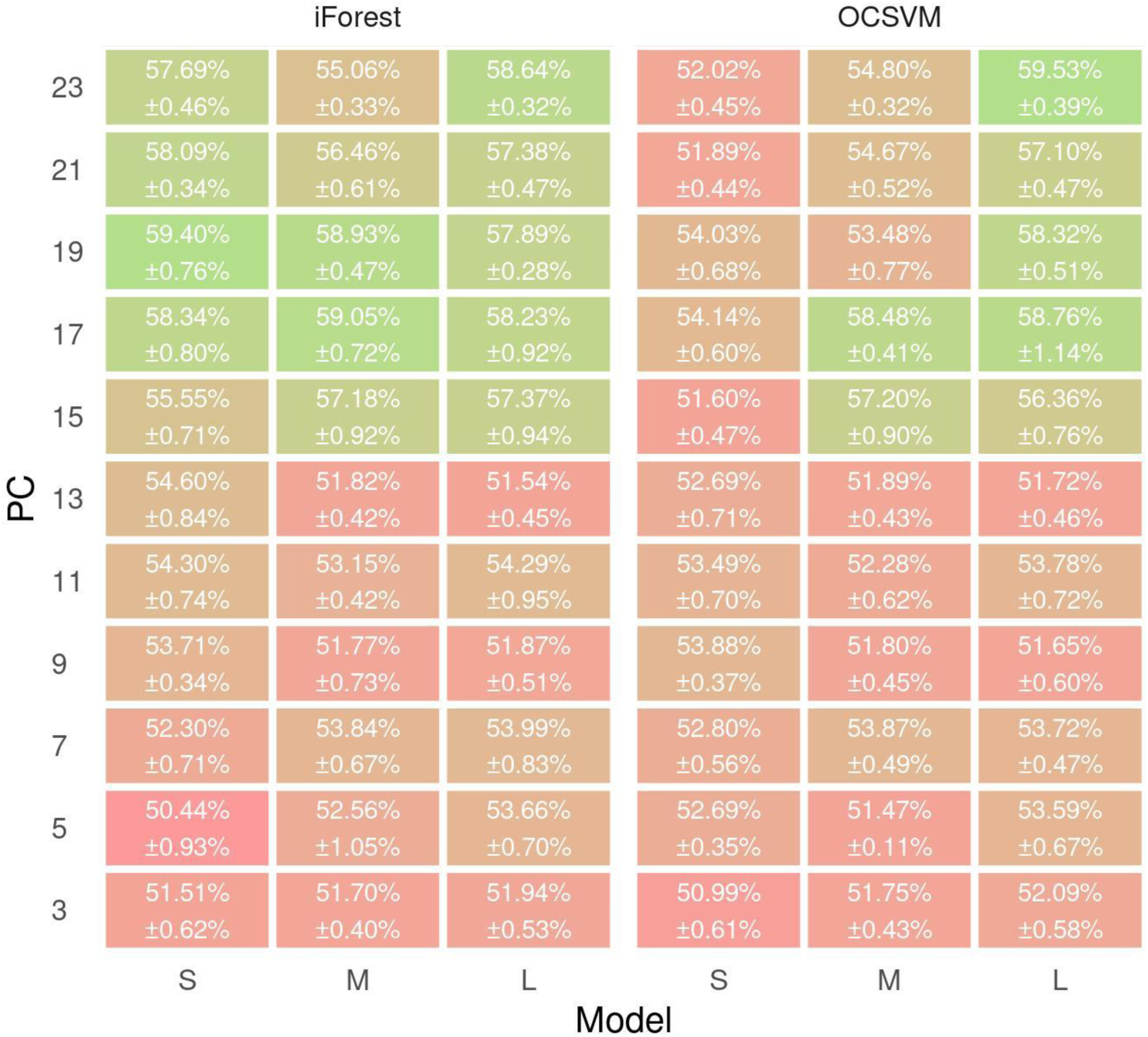
The precision metrics with standard errors for S, M, and L model architectures across different numbers of fitted principal components (PC).

Based on the precision metric and MAE, the M model that fits 17 PCs and the iForest classification were selected as the best procedure and were further used in all downstream analyses. Figure 7. presents the density of the MAE across PCs for each SNP underlying the four classification groups - TP, TN, FP, and FN for two algorithms - iForest and OCSVM with the intersection of the TP class. It clearly indicates that SNPs classified as incorrect (TP and FP) were associated with higher reconstruction errors, with the highest MAEs corresponding to systematically incorrectly called SNPs labelled as TP. Still, there was considerable overlap between densities underlying TP, TN, FP, and FN classes. Moreover, the variability of MAE of the incorrectly called SNPs is higher than that of the correctly called SNPs.

**Figure 7.**
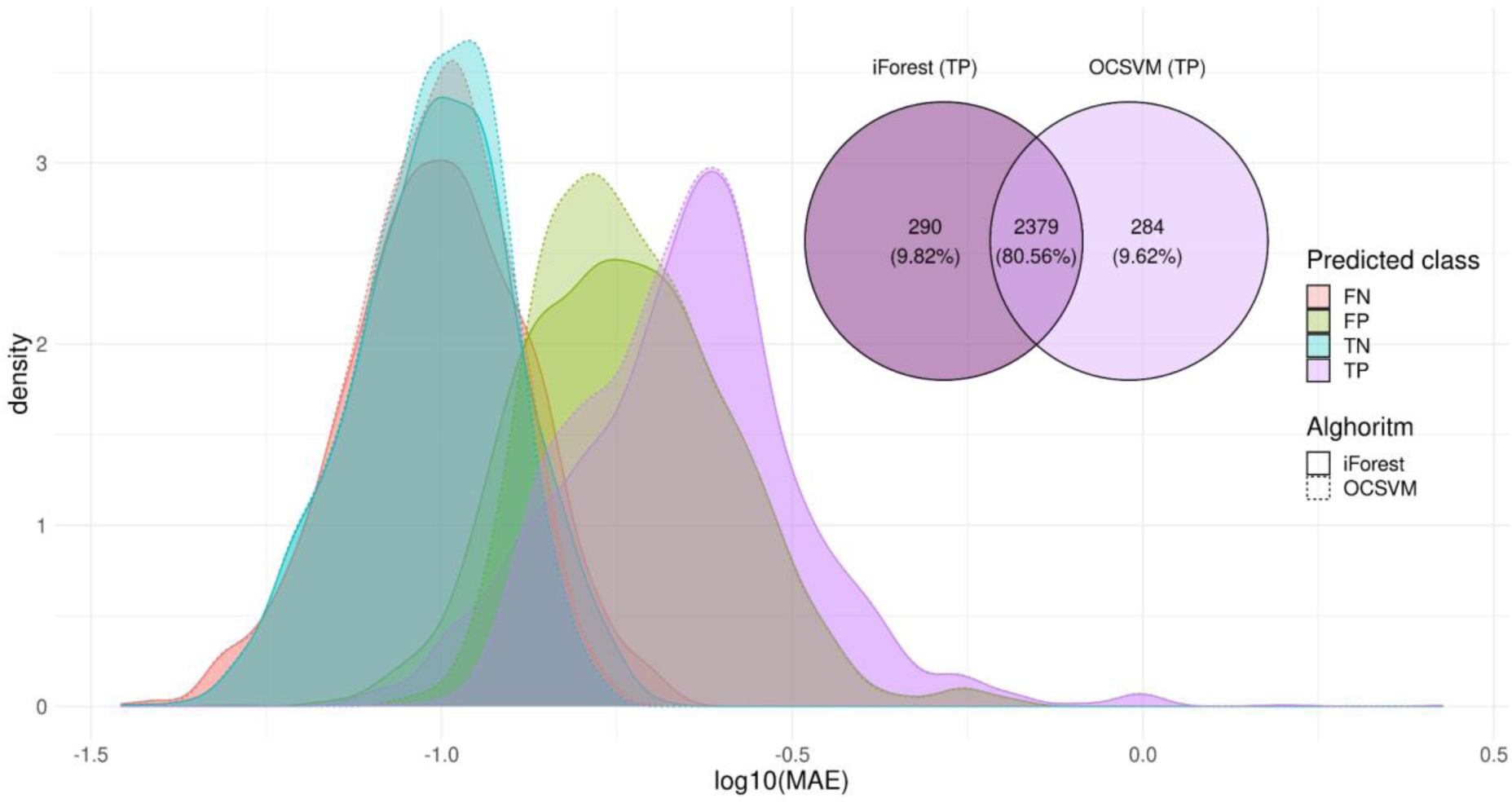
The density of MAE across PCs for each SNP classification group underlying model M that fits 17 principal components followed by the classification based in iForest and OCSVM with the intersection of the TP class between two algorithms.

### Frequency distribution of nucleotides surrounding incorrectly called SNPs

To investigate the influence of nucleotide composition on an incorrect SNP call, we examined its surrounding genomic context. Figures 8 and 9. show the frequency of nucleotides at four positions downstream (Figure 8) and upstream (Figure 9) of an incorrect SNP call, calculated separately for every possible incorrect substitution. No general pattern of nucleotide frequencies near an incorrect call was identified. However, for the erroneous substitution of A by T, there was an excess of thymine at a neighbouring downstream position and at the 2nd upstream position that may have influenced the fluorescent labelling of the nucleotide reading. Furthermore, for a wrong call of G in place of C, an excess of guanine in the second upstream position could be observed.

**Figure 8.**
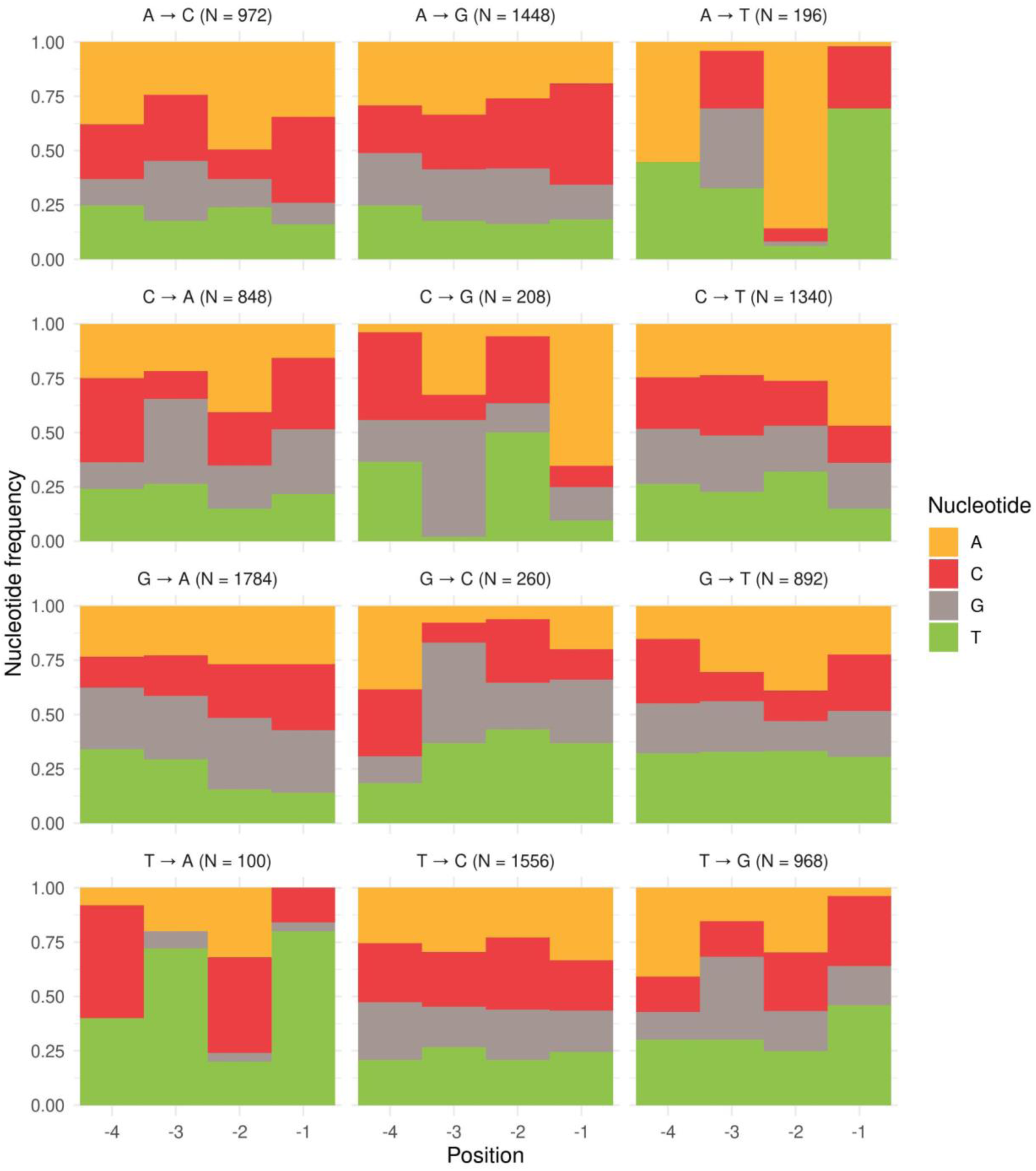
The frequency of nucleotides at the four positions (-1 to -4) downstream of an incorrectly called SNP from the TP class, shown separately for each incorrect substitution with the corresponding number (N) of occurrences. Nucleotide colour coding reflects the fluorescent labelling used by the Illumina NovaSeq 6000 sequencer.

**Figure 9.**
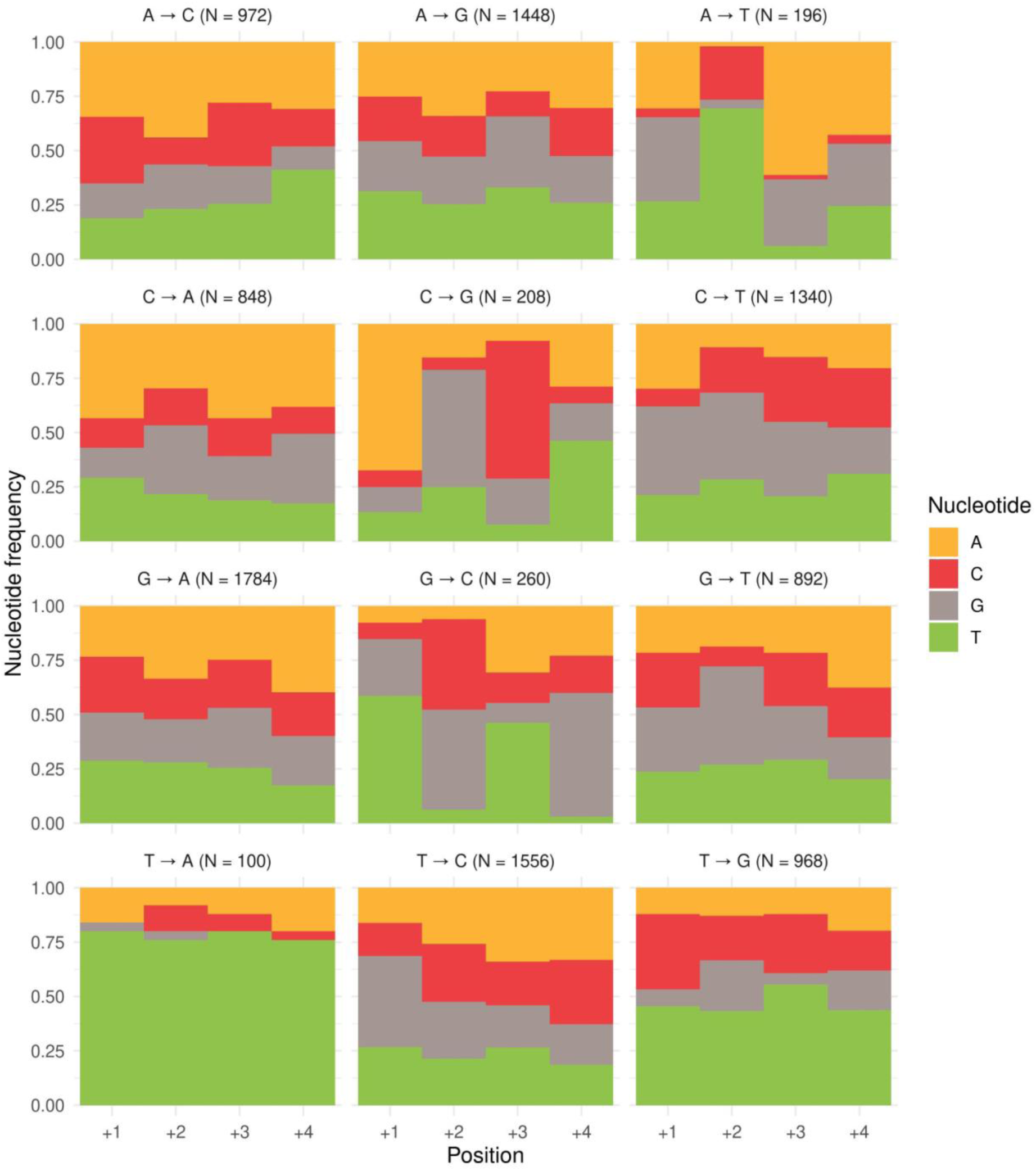
The frequency of nucleotides at the four positions (+1 to +4) upstream of an incorrectly called SNP from the TP class, shown separately for each incorrect substitution with the corresponding number (N) of occurrences. Nucleotide colour coding reflects the fluorescent labelling used by the Illumina NovaSeq 6000 sequencer.

In addition, we also considered trimer motif frequencies at incorrectly called SNP sites. A trimer was defined as three base pairs leading up to and including the incorrectly called base were selected to examine the triplets associated with the systematic error rate, which was expressed by the ratio of a triplet frequency within the TP class (corresponding to systematically incorrect SNP calls) and a triplet frequency in the whole test dataset. Hence, the ratio of one expresses no systematic effect of a given triplet on the incorrect SNP call. Figure 10. shows the frequency with a corresponding 95% confidence interval of ten triplets out of 64 possible motifs with the highest frequency. The most error-prone triplet was CGC. In general, 28 trimers have a frequency above 1.0, indicating that they are more likely to be associated with errors.

**Figure 10.**
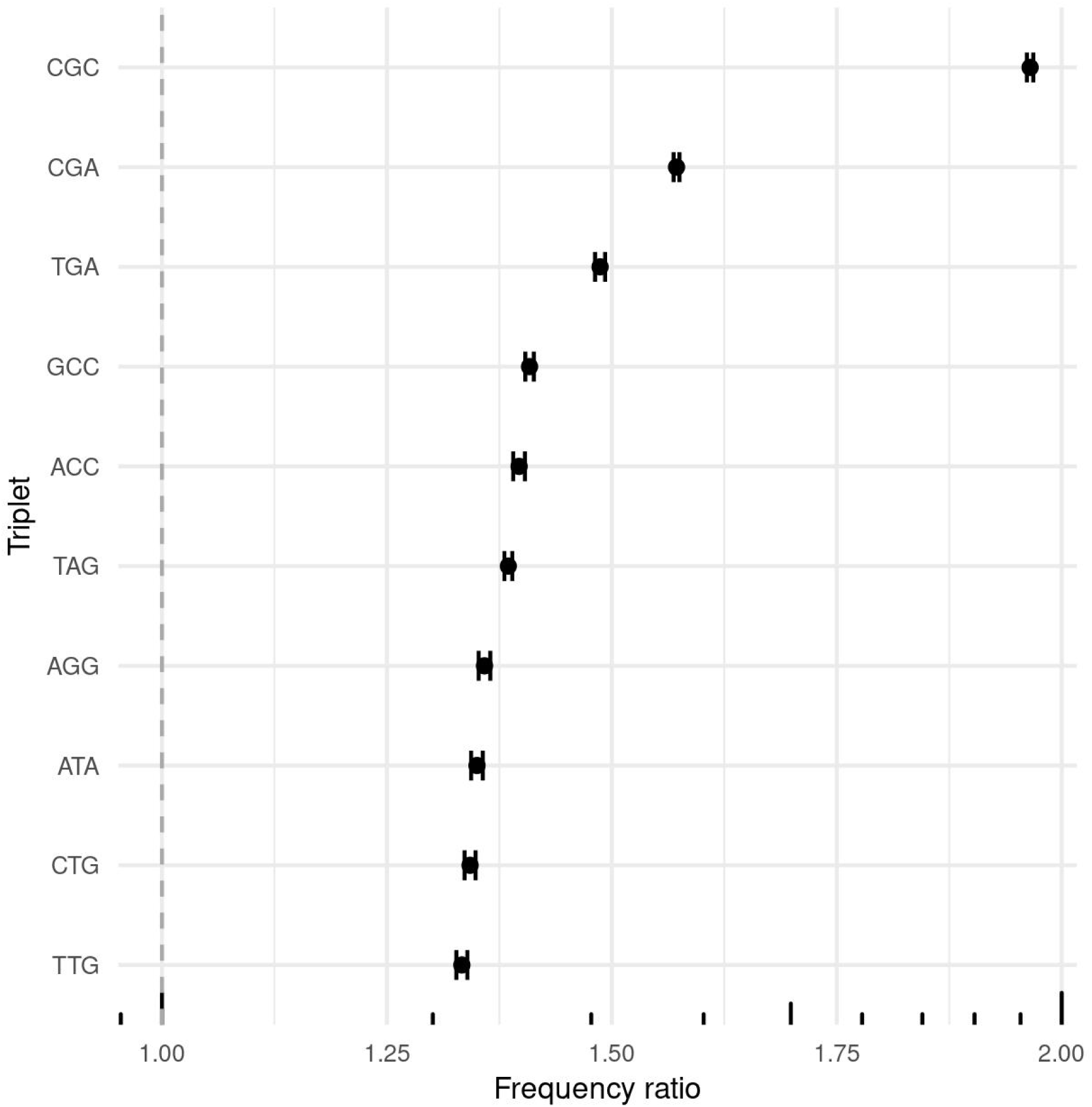
The frequency of the ten most frequent triplet sequences leading up to and including the incorrectly called site with their 95% confidence intervals.

## DISCUSSION

### Identification of systematically incorrectly called SNPs

Converting raw DNA sequence data to SNP genotype calls involves several computational steps. Typically output is a VCF file containing *predictions* of true SNP genotypes, each of them associated with some non-unity accuracy. Unfortunately, in the VCF file the accuracy is not expressed by a single value, since no formal estimation model exists. Instead there are several indicators of the probability of the called SNP representing the true genotype, such as DP or GQ metrics (30, 31). The influence of the nucleotide context surrounding the called SNP on the probability of calling the true SNP genotype has been investigated (32). However, this information is not available in standard VCF files. However, strategies have been proposed to take nucleotide context into account during SNP calling. For example, GATK’s HaplotypeCaller, widely used for variant detection, implements a Bayesian model to estimate the likelihood of each possible haplotype and genotype, considering the nucleotide context around each SNP (21). Although the accuracy of SNP calling has improved over time (33), our results demonstrated that even modern sequencers, such as the Illumina NovaSeq 6000, still deliver incorrectly identified SNP genotypes originating from WGS data.

In our analysis, we focussed on two aspects of the identification of incorrectly called SNPs. First, we investigated a *technical aspect* of the architecture underlying the DL-based classifier of correctly and incorrectly called SNPs that was used to differentiate between systematically incorrectly called genotypes that are due to specific constellations of explanatory variables and random errors in called genotypes. Then, only for systematic errors, a *biological aspect* of the influence of the specific sequence context surrounding the polymorphic site on calling the incorrect genotype was considered. Our analysis demonstrated that 59.53% (±0.39%) of the incorrectly called SNPs can be explained by the systematic pattern of explanatory variables, while the remainder represents random errors not associated with explanatory variables available in the standard VCF output nor related to sequence context.

### Exploring autoencoder architectures and data representations

Our previous work on the classification of correctly and incorrectly called SNPs from a similar data set (34), employed a traditional DL-based classifier incorporating a set of dense neural networks yielded moderate success in classification quality, was largely due to the extreme class count imbalance between the correctly and incorrectly called SNPs. To address class imbalance problem, we adapted a different approach by fitting DL-based autoencoders to the set of correctly called SNPs only and then performing the classification on this already trained algorithm. Autoencoders have been widely used for the detection of anomalies since they can efficiently capture complex patterns of explanatory variables characteristic of correct (i.e. non-anomalous) outcomes (35, 36). However, to our knowledge, no autoencoder-based classifier of correctly and incorrectly called SNPs has been proposed for WGS data. Therefore, we created an optimised autoencoder architecture. This was done by exploring various activation functions (tanh, sigmoid, ReLU), various model architectures (S, M, L), and various ways of data representation (no transformation and various numbers of PCs). The predictive performance of the autoencoders did not vary considerably among configurations. The most profound impact on classification quality was attributed to the interplay between model architectures and the number of principal components. In particular, the contribution of the explanatory variables to the data representation revealed a clear pattern. Features extracted from a standard VCF file, especially ALT, REF, GT:A, and GT:B have the largest loadings on the first three PCs, indicating their important role in the explanation of data quality. However, the ’fine tuning’ of data differentiation is clearly due to the sequence context expressed by the four nucleotides downstream and upstream of the call, which was expressed by the higher contribution of the sequence context to higher PCs (> 3) PCs. This pattern can also be observed on the classification precision heatmap (Figure 6). Here, precision increased after capturing more of data variability by considering higher dimensions expressed by PCs > 3. However, a precision plateau occurred after the model dimensionality reached 17 PCs. In none of the tested architectures, higher dimensionalities increased classification precision. The precision of 59.53% (±0.39%) was obtained for the best architecture of the M model, with the ELU activation function with 17 PCs.

### Impact of sequence context on systematically incorrectly called SNPs

The influence of sequence context on systematically incorrectly called SNPs is an important topic in the field of genomics and genetic analysis (37, 38). The studies of Bansal et al. (39) and Meacham et al. (37) provide valuable insights into the differentiation of systematic error and random SNP calling errors, with a focus on the sequence context surrounding them. Bansal et al. (2010) defined systematic errors as incorrectly called SNP sites that are consistent across multiple individuals, indicating a potential underlying systematic effect in either the sequencing or the SNP calling process. In contrast, Meacham et al (2011) defined systematic base calling errors by using a logistic regression classifier. Our approach to defining the systematic base calling errors differs from those adapted in the aforementioned studies and is based on the anomaly detection paradigm underlying the autoencoder model. The observed agreement between outlier detection methods - iForest and OCSVM, serves as validation of the definition of systematically incorrect SNP calls adopted in our study and underscores the presence of a systematic pattern underlying these errors.

The sequence context surrounding systematic error sites identified in other studies varies among published reports and in many cases differs from our findings. Various reasons contribute to this disagreement. First, different studies adapt different definitions of systematic errors (32, 37, 39–41), while some other studies do not differentiate between systematic and random incorrect calls (4, 42). Second, different studies analysed data generated with different sequencing technologies, that might differ regarding sequence context error profiles. Our results are valid for the sequencing methodology underlying the Illumina NovaSeq 6000 sequencer. The influence of nucleotides directly preceding the systematically misidentified SNPs, accurately reflects the chemistry underlying fluorescent nucleotide imaging utilized in the Illumina NovaSeq 6000 sequencer. Specifically, the distribution of fluorescent signals corresponding to the four nucleotides complicates differentiation between A and C, as well as between A and T. This difficulty is mirrored in the pattern of incorrect substitutions identified for systematically misidentified SNPs in our dataset (Figure 8). Our findings demonstrate that A/T errors are correlated with the presence of T, at the first downstream position, indicating that the mislabelling of the SNP may result from the similar fluorescence pattern of the nucleotide directly preceding it. Understanding the dynamics of error occurrence within DNA sequences can also be achieved by examining the frequency of trimer motifs at error sites (40). The high frequency of the CGC trimer can be explained by mistaking a C call. This mistake might occur because the NovaSeq 6000 sequencing platform interprets a lack of fluorescence as the nucleotide G. Consequently, it may be possible to incorrectly identify C within the CGC trimer if fluorescence is found at the -2 downstream position. Some of the motifs such as CGA were also reported in another study (4) or GCC (40). Also, there is no consistency in the literature regarding the frequency of particular nucleotide substitutions constituting systematic wrong SNP calls. However, for the NovaSeq 6000, a close agreement between our results and the results reported by Ma et al. (42) points to A/G and T/C substitutions having the highest error rates.

In our investigation, we focussed on two crucial elements in identifying incorrectly called SNPs. First, we conducted an analysis of the architecture underlying the AI-based anomaly detection approach implemented as an AE-classifier that was used to differentiate between correct and incorrect genotype calls based on the combination by PCA of standard explanatory variables from a VCF with additional four reference nucleotides upstream and four downstream, representing the sequence context surrounding the SNP site. This technical aspect explained the complex details of the classifier’s functioning and its ability to differentiate between different genotype-calling errors, as a result providing valuable insights into the differentiation of systematic error and random SNP calling errors. Next, we focus on a biological exploration of the impact that sequence context has on the polymorphic nucleotide call. We were able to find important patterns between the sequence context and the tendency to identify the wrong genotypes. In general, our examination that integrates insights from both domains will improve the repeatability and reliability of SNP calling itself as well as of downstream genetic analyses, by the possibility of the differential weighting of the SNP genotypes estimated by SNP calling pipelines, depending not only on the explanatory variables available in the VCF file but also on the neighbouring sequence composition. Finally, we demonstrated that, since biological data typically associate with various constraints of the financial, ethical, or sampling nature, it is often impossible to obtain very large samples. In such a situation, applications involving machine learning approaches that originally have been developed for very large and informative data structures, need to be fine-tuned in terms of selecting the best models’ architectures, their hyperparameters as well as best input representations. In the case of a limited size of input data, only such an extensive model selection approach allows for meaningful statistical inferences.

## DATA AVAILABILITY

Deoxyribonucleic acid sequences of 20 cows from the training data set are available from the NCBI BioProject database under the accession ID PRJNA979229.

## AUTHOR CONTRIBUTIONS

Krzysztof Kotlarz: Conceptualization, Data curation, Formal analysis, Methodology, Validation, Writing – original draft. Magda Mielczarek: Data curation, Formal Analysis, Writing—review & editing. Przemysław Biecek: Methodology, Supervision. Bernt Guldbrandtsen: Methodology, Writing—review & editing. Joanna Szyda: Funding acquisition, Investigation, Project administration, Supervision, Writing – original draft.

## ACKNOWLEDGEMENTS

Calculations were carried out at the Wroclaw Centre for Networking and Supercomputing and Poznan Supercomputing and Networking Center.

## REFERENCES

1. Koboldt, D.C., Steinberg, K.M., Larson, D.E., Wilson, R.K. and Mardis, E.R. (2013) The Next-Generation Sequencing Revolution and Its Impact on Genomics. Cell, 155, 27–38.

2. Hartfield, M., Poulsen, N.A., Guldbrandtsen, B. and Bataillon, T. (2021) Using singleton densities to detect recent selection in *Bos taurus*. Evol Lett, 5, 595–606.

3. Olson, N.D., Lund, S.P., Colman, R.E., Foster, J.T., Sahl, J.W., Schupp, J.M., Keim, P., Morrow, J.B., Salit, M.L. and Zook, J.M. (2015) Best practices for evaluating single nucleotide variant calling methods for microbial genomics. Front Genet, 6, 235.

4. Stoler, N. and Nekrutenko, A. (2021) Sequencing error profiles of Illumina sequencing instruments. NAR Genom Bioinform, 3, lqab019.

5. Fox, E.J., Reid-Bayliss, K.S., Emond, M.J. and Loeb, L.A. (2014) Accuracy of Next Generation Sequencing Platforms. Next Gener Seq Appl, 1, 1000106.

6. Krachunov, M., Nisheva, M. and Vassilev, D. (2017) Machine learning models in error and variant detection in high-variation high-throughput sequencing datasets. Procedia Comput Sci, 108, 1145–1154.

7. King, G. and Zeng, L. (2001) Logistic Regression in Rare Events Data. Political Analysis, 9, 137–163.

8. King, G. and Zeng, L. (2001) Explaining Rare Events in International Relations. Int Organ, 55, 693–715.

9. Wang, H., Xu, Q. and Zhou, L. (2015) Large Unbalanced Credit Scoring Using Lasso-Logistic Regression Ensemble. PLoS One, 10, e0117844.

10. Zhang, L., Geisler, T., Ray, H. and Xie, Y. (2022) Improving logistic regression on the imbalanced data by a novel penalized log-likelihood function. J Appl Stat, 49, 3257– 3277.

11. Frühwirth-Schnatter, S. and Wagner, H. (2008) Marginal likelihoods for non-Gaussian models using auxiliary mixture sampling. Comput Stat Data Anal, 52, 4608–4624.

12. Sakurada, M. and Yairi, T. (2014) Anomaly Detection Using Autoencoders with Nonlinear Dimensionality Reduction. In Proceedings of the MLSDA 2014 2nd Workshop on Machine Learning for Sensory Data Analysis. ACM, New York, NY, USA, 4–11.

13. Zavrak, S. and Iskefiyeli, M. (2020) Anomaly-Based Intrusion Detection From Network Flow Features Using Variational Autoencoder. IEEE Access, 8, 108346–108358.

14. Torabi, H., Mirtaheri, S.L. and Greco, S. (2023) Practical autoencoder based anomaly detection by using vector reconstruction error. Cybersecurity, 6, 1.

15. Andrews, S. (2010) FastQC: A Quality Control Tool for High Throughput Sequence Data.

16. Ewels, P., Magnusson, M., Lundin, S. and Käller, M. (2016) MultiQC: summarize analysis results for multiple tools and samples in a single report. Bioinformatics, 32, 3047–3048.

17. Bolger, A.M., Lohse, M. and Usadel, B. (2014) Trimmomatic: a flexible trimmer for Illumina sequence data. Bioinformatics, 30, 2114–2120.

18. Li, H. and Durbin, R. (2009) Fast and accurate short read alignment with Burrows– Wheeler transform. Bioinformatics, 25, 1754–1760.

19. Li, H., Handsaker, B., Wysoker, A., Fennell, T., Ruan, J., Homer, N., Marth, G., Abecasis, G. and Durbin, R. (2009) The Sequence Alignment/Map format and SAMtools. Bioinformatics, 25, 2078–2079.

20. Quinlan, A.R. and Hall, I.M. (2010) BEDTools: a flexible suite of utilities for comparing genomic features. Bioinformatics, 26, 841–842.

21. McKenna, A., Hanna, M., Banks, E., Sivachenko, A., Cibulskis, K., Kernytsky, A., Garimella, K., Altshuler, D., Gabriel, S., Daly, M., et al. (2010) The Genome Analysis Toolkit: A MapReduce framework for analyzing next-generation DNA sequencing data. Genome Res, 20, 1297–1303.

22. Lê, S., Josse, J. and François Husson (2008) FactoMineR: A Package for Multivariate Analysis. J Stat Softw, 25, 1–18.

23. Chollet, F. and others (2015) Keras.

24. Abadi, M., Barham, P., Chen, J., Chen, Z., Davis, A., Dean, J., Devin, M., Ghemawat, S., Irving, G., Isard, M., et al. (2016) TensorFlow: A system for large-scale machine learning. In 12th USENIX Symposium on Operating Systems Design and Implementation 265–283.

25. Clevert, D.-A., Unterthiner, T. and Hochreiter, S. (2015) Fast and Accurate Deep Network Learning by Exponential Linear Units (ELUs), arXiv doi: 1511.07289.

26. Kingma, D.P. and Ba, J. (2014) Adam: A Method for Stochastic Optimization, arXiv:1412.6980v9.

27. Pedregosa, F., Varoquaux, G., Gramfort, A., Michel V. and Thirion, B., Grisel, O., Blondel, M., Prettenhofer P. and Weiss, R., Dubourg, V., Vanderplas, J., Passos, A., et al. (2011) Scikit-learn: Machine Learning in Python. Journal of Machine Learning Research, 12, 2825–2830.

28. Amer, M., Goldstein, M. and Abdennadher, S. (2013) Enhancing one-class support vector machines for unsupervised anomaly detection. In *Proceedings of the ACM SIGKDD Workshop on Outlier Detection and Description*. ACM, New York, NY, USA, 8–15.

29. Liu, F.T., Ting, K.M. and Zhou, Z.-H. (2008) Isolation Forest. In 2008 Eighth IEEE International Conference on Data Mining. IEEE, 413–422.

30. Zook, J.M., McDaniel, J., Olson, N.D., Wagner, J., Parikh, H., Heaton, H., Irvine, S.A., Trigg, L., Truty, R., McLean, C.Y., et al. (2019) An open resource for accurately benchmarking small variant and reference calls. Nat Biotechnol, 37, 561–566.

31. Krusche, P., Trigg, L., Boutros, P.C., Mason, C.E., De La Vega, F.M., Moore, B.L., Gonzalez-Porta, M., Eberle, M.A., Tezak, Z., Lababidi, S., et al. (2019) Best practices for benchmarking germline small-variant calls in human genomes. Nat Biotechnol, 37, 555–560.

32. Dohm, J.C., Lottaz, C., Borodina, T. and Himmelbauer, H. (2008) Substantial biases in ultra-short read data sets from high-throughput DNA sequencing. Nucleic Acids Res, 36, e105.

33. Jasper, R.J., McDonald, T.K., Singh, P., Lu, M., Rougeux, C., Lind, B.M. and Yeaman, S. (2022) Evaluating the accuracy of variant calling methods using the frequency of parent-offspring genotype mismatch. Mol Ecol Resour, 22, 2524–2533.

34. Kotlarz, K., Mielczarek, M., Suchocki, T., Czech, B., Guldbrandtsen, B. and Szyda, J. (2020) The application of deep learning for the classification of correct and incorrect SNP genotypes from whole-genome DNA sequencing pipelines. J Appl Genet, 61, 607–616.

35. Chen, J., Sathe, S., Aggarwal, C. and Turaga, D. (2017) Outlier Detection with Autoencoder Ensembles. In Proceedings of the 2017 SIAM International Conference on Data Mining. Society for Industrial and Applied Mathematics, Philadelphia, PA, 90–98.

36. Zhou, C. and Paffenroth, R.C. (2017) Anomaly Detection with Robust Deep Autoencoders. In Proceedings of the 23rd ACM SIGKDD International Conference on Knowledge Discovery and Data Mining. ACM, New York, NY, USA, 665–674.

37. Meacham, F., Boffelli, D., Dhahbi, J., Martin, D.I., Singer, M. and Pachter, L. (2011) Identification and correction of systematic error in high-throughput sequence data. BMC Bioinformatics, 12, 451.

38. Taub, M.A., Corrada Bravo, H. and Irizarry, R.A. (2010) Overcoming bias and systematic errors in next generation sequencing data. Genome Med, 2, 87.

39. Bansal, V., Harismendy, O., Tewhey, R., Murray, S.S., Schork, N.J., Topol, E.J. and Frazer, K.A. (2010) Accurate detection and genotyping of SNPs utilizing population sequencing data. Genome Res, 20, 537–545.

40. Nakamura, K., Oshima, T., Morimoto, T., Ikeda, S., Yoshikawa, H., Shiwa, Y., Ishikawa, S., Linak, M.C., Hirai, A., Takahashi, H., et al. (2011) Sequence-specific error profile of Illumina sequencers. Nucleic Acids Res, 39, e90–e90.

41. Allhoff, M., Schönhuth, A., Martin, M., Costa, I.G., Rahmann, S. and Marschall, T. (2013) Discovering motifs that induce sequencing errors. BMC Bioinformatics, 14, S1.

42. Ma, X., Shao, Y., Tian, L., Flasch, D.A., Mulder, H.L., Edmonson, M.N., Liu, Y., Chen, X., Newman, S., Nakitandwe, J., et al. (2019) Analysis of error profiles in deep next-generation sequencing data. Genome Biol, 20, 50.

